# ‘Silent Spring’ revisited: sublethal neonicotinoid exposure suppresses birdsong

**DOI:** 10.64898/2026.11.16.766774

**Authors:** Ednei B. dos Santos, Jaemin Lee, Sang-im Lee, Satoshi Kojima

## Abstract

Neonicotinoids are widely used pesticides linked to impaired reproduction and population decline in birds. However, the behavioral mechanisms connecting exposure to reduced reproduction remain unclear. Here, we examined the effects of imidacloprid, one of the most commonly used neonicotinoids, on singing in male zebra finches. Birds received sublethal doses of imidacloprid through drinking water over 30 days, followed by a 30-day washout. Using continuous acoustic monitoring and female-presentation assays, we quantified non-courtship and female-directed courtship song and assessed body mass, locomotion, non-vocal displays, and song structure. During the 30-day drinking-water exposure, imidacloprid reduced daily non-courtship singing, with effects persisting into washout. It also reduced courtship song, whereas body mass, locomotion, non-vocal displays, and song structure showed no detectable treatment-related changes. In a separate experiment, acute imidacloprid injection also reduced courtship song but was accompanied by some non-vocal behavioral impairment. These findings identify song output as a sensitive behavioral endpoint for neonicotinoid toxicity, suggesting that even low doses might impair songbird reproduction by suppressing singing before conventional signs of toxicity become apparent. This suppression of song production implies that widespread neonicotinoid use could silently contribute to population declines in wild birds by compromising their ability to attract mates, defend territories, form pair bonds, and successfully breed. Song output may therefore complement conventional measures when assessing the sublethal effects of neonicotinoids in songbirds.

## Introduction

As Rachel Carson famously warned in *Silent Spring* [1], the indiscriminate use of chemical insecticides can severely impact natural environments [2] and contribute to declines in wildlife populations, including birds. More recently, neonicotinoid insecticides, which are widely used in agriculture, livestock production, and gardening [3–5], have been implicated in avian population declines by numerous studies. For example, in the Netherlands, surface-water neonicotinoid concentrations were associated with long-term declines in insectivorous birds [6], while county-level neonicotinoid use in the United States was associated with reduced abundance and diversity, especially among grassland and insectivorous species [7]. These effects on birds are thought to arise not only from reduced availability of insect prey [8] but also from direct exposure through treated seeds, contaminated plant material, and drinking water. Indirect exposure through contaminated arthropod prey has also been suggested [9,10]. However, the mechanisms that connect such exposure to reduced survival and reproduction remain poorly understood.

Evidence from experimental studies has accumulated rapidly, showing that neonicotinoid exposure adversely affects avian physiology, reproduction, survival, and behavior [11]. For example, in white-crowned sparrows, neonicotinoid exposure reduced migratory fueling and delayed migration [12]. Similarly, studies have reported reduced energy expenditure in ruby-throated hummingbirds [13], reduced sperm density in house sparrows [14], disruption of the pituitary–thyroid axis in red munias [15], and reduced breeding investment and offspring immunity in red-legged partridges [16]. These findings indicate that neonicotinoids can disturb metabolic, neuroendocrine, and reproductive processes of birds even at sublethal doses. Yet the behaviors through which exposure may reduce reproductive opportunities have received comparatively little attention. One such possibility is that pesticides may silence birds not only by reducing their populations, as depicted in *Silent Spring*, but also by suppressing singing in birds that survive exposure. To our knowledge, only one previous study has reported reduced singing following imidacloprid exposure, but singing was scored at the group level rather than measured quantitatively in individual birds [17].

Songbirds use song to attract mates, defend territories, establish and maintain pair bonds, and coordinate social interactions [18]. Reductions in song rate or quality could therefore substantially influence access to mates and reproductive opportunities, even when birds remain outwardly active. Singing also serves as vocal practice. High rates of song production are required for juveniles to acquire mature song [19,20], and continued practice is needed to maintain optimal song performance in adulthood [21,22]. A reduction in singing could thus affect not only immediate communication but also the development and maintenance of song performance that may influence reproductive success. The possibility that neonicotinoids affect singing behavior is also supported by the close link between their mode of action and the neural mechanisms underlying song. Neonicotinoids were developed to act on insect nicotinic acetylcholine receptors (nAChRs). They can also interact with vertebrate nAChRs, which are abundantly expressed in the brains of birds and other vertebrates, and perturb neural and physiological processes [23,24]. Because, in songbirds, nAChRs are expressed throughout nuclei involved in vocal perception, learning, and production [25–28], neonicotinoids could act through these receptors to alter the neural processes that initiate and sustain singing, without necessarily changing gross locomotion or the acoustic structure of the songs produced. Nevertheless, to our knowledge, no study has quantitatively tested how neonicotinoid exposure affects daily song output or female-directed courtship singing in songbirds.

To address this gap, we investigated the effects of sublethal exposure to imidacloprid, a widely used neonicotinoid insecticide, on singing behavior in adult male zebra finches (*Taeniopygia guttata*), a well-established model for vocal learning and production [29]. Specifically, we continuously monitored non-courtship song (commonly referred to as “undirected song”) through a 30-day drinking-water exposure and a subsequent 30-day washout. We combined this monitoring with female-presentation assays to quantify female-directed courtship song. We also measured body mass, locomotion, non-vocal courtship behavior, and song structure to determine whether any change in song output reflected broader behavioral impairment or impaired vocal performance. Finally, in a within-subject experiment, we examined the immediate effects of acute imidacloprid injection on courtship song and associated courtship behaviors to assess how the behavioral effects of drinking-water exposure and acute exposure may differ.

## Results

### Imidacloprid exposure through drinking water reduces daily undirected singing

To determine whether imidacloprid exposure through drinking water altered singing behavior, we quantified daily undirected song bouts across exposure phases and washout (**Fig 1A**). Imidacloprid exposure through drinking water reduced daily undirected singing across successive doses. A negative-binomial generalized linear mixed model, with bird identity as a random effect, detected a treatment-by-experimental phase interaction for full-day song output (χ²₄ = 35.17, *P* < 0.0001). Contrasts between each phase and its corresponding baseline showed that treated males sang less during the initial phase at 1% of the estimated LD₅₀ (**Fig 1B,C; S1 Table**). This reduction persisted at 2.5% and 5% of the estimated LD₅₀, with the largest effect observed at 5%, and remained detectable during the final 10 days of the 30-day washout. A similar pattern was observed for songs produced during the first 2 h after lights-on, the time of day when birds generally sing most intensely (χ²₄ = 41.67, *P* < 0.0001; **S1 Fig, panels A and B**). An exploratory cumulative analysis, repeated after each exposure day, first detected a treatment difference after three days for full-day song output (χ²₁ = 4.75, *P* = 0.029; **Fig 1C**) and after seven days for the first 2 h after lights-on (χ²₁ = 5.46, *P* = 0.019; **S1 Fig, panel A**). Thus, imidacloprid reduced daily song output across the full light period and during peak morning singing, with effects persisting into late washout. Body mass did not show treatment-specific changes across exposure or washout (**Fig 1D; S2 Table)**, and no overt lethargy, postural impairment or loss of balance was observed during drinking-water exposure.

**Fig 1.**
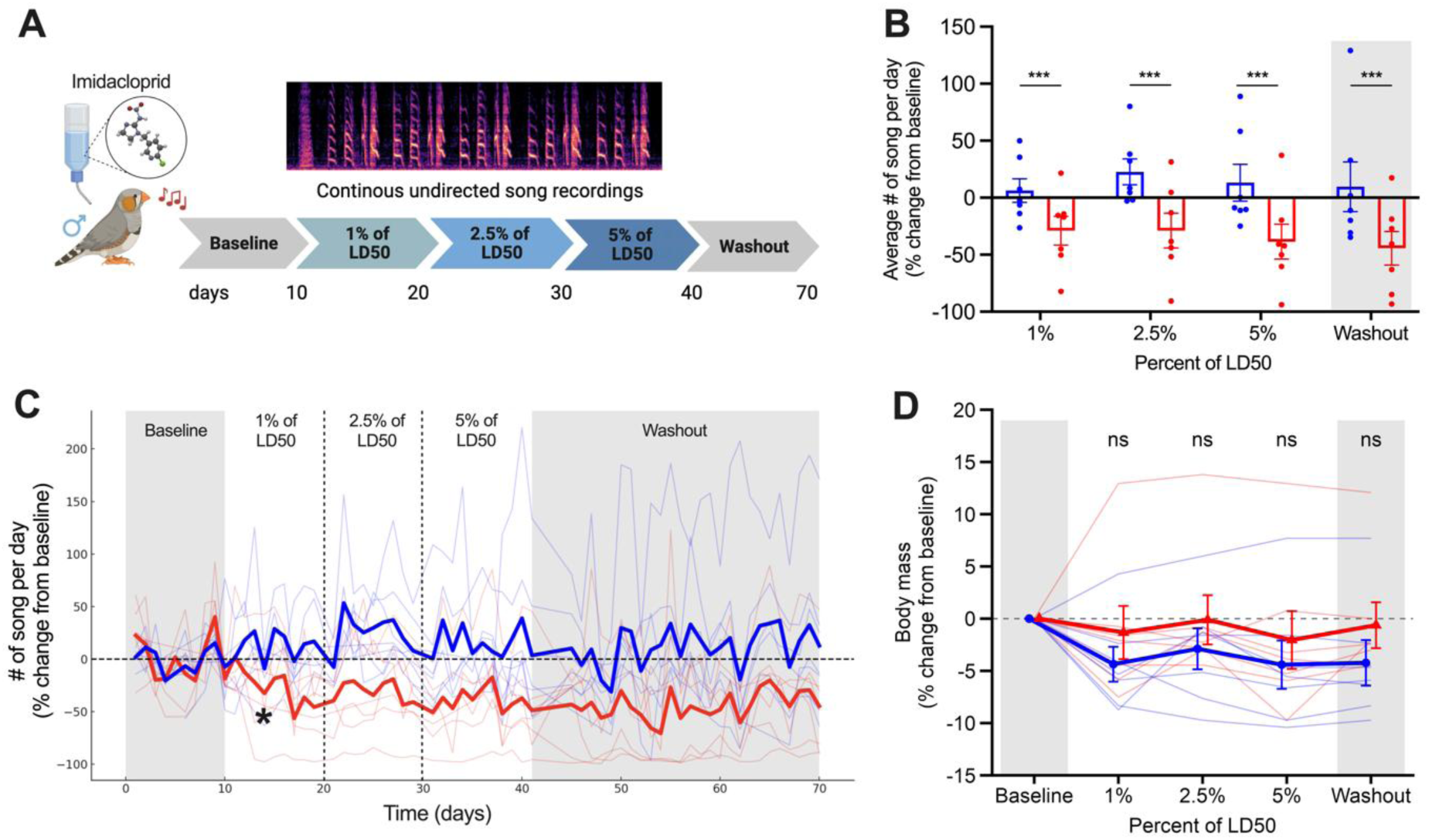
Imidacloprid exposure through drinking water reduces undirected singing without detectable changes in body mass. **(A)** Experimental design: adult males were recorded continuously during a 10-day baseline, three successive 10-day drinking-water exposure phases (1%, 2.5% and 5% of the estimated LD₅₀) and a 30-day washout. **(B)** Mean full-day song output during each exposure phase and late washout (% change from baseline); points, individual males; bars, group mean ± s.e.m. Full-day counts were analyzed with a negative-binomial generalized linear mixed model (treatment × experimental phase interaction: χ²₄ = 35.17, *P* < 0.0001). **(C)** Daily number of undirected song bouts (% change from each male’s baseline); thin lines, individual males; thick lines, group means. The asterisk marks the first exposure day on which a statistically significant difference between control and treated males was detected (third day of exposure; χ²₁ = 4.75, *P* = 0.029), with the analysis repeated using all data collected up to each exposure day. **(D)** Body mass across timepoints (% change from baseline); thin lines, individual males; thick lines, group means ± s.e.m. Body mass was analyzed with a linear mixed model (no treatment-specific change; all Holm-adjusted *P* ≥ 0.43). Control, blue; imidacloprid-treated, red. *n* = 7 control and 7 imidacloprid-treated males. ns, not significant; \*\*\**P* < 0.001.

### Imidacloprid exposure through drinking water suppresses courtship song

We next tested whether imidacloprid reduced female-directed courtship singing. After drinking-water exposure, males underwent four counterbalanced 5-min female-presentation sessions across two days (**Fig 2A**). Treated males produced fewer song bouts than controls (χ²₁ = 10.14, *P* = 0.0014; **Fig 2B; S3 Table**), and motif counts showed the same pattern (χ²₁ = 10.99, *P* = 0.00091; **Fig 2C**). To determine whether this reflected a selective effect on singing or broader suppression of courtship activity, we quantified female approach, locomotion and non-vocal displays from video recordings. Treated and control males spent similar proportions of time adjacent to the female cage (χ²₁ = 0.003, adjusted *P* = 0.96; **Fig 2E**) and did not differ in zone transitions (χ²₁ = 0.14, adjusted *P* = 0.96; **Fig 2F**), perch changes (χ²₁ = 0.034, adjusted *P* = 0.96; **Fig 2G**) or bill wipes (χ²₁ = 0.96, adjusted *P* = 0.96; **Fig 2H**). Hence, courtship song suppression occurred without detectable reductions in locomotor activity, female approach or the non-vocal behaviors measured.

**Fig 2.**
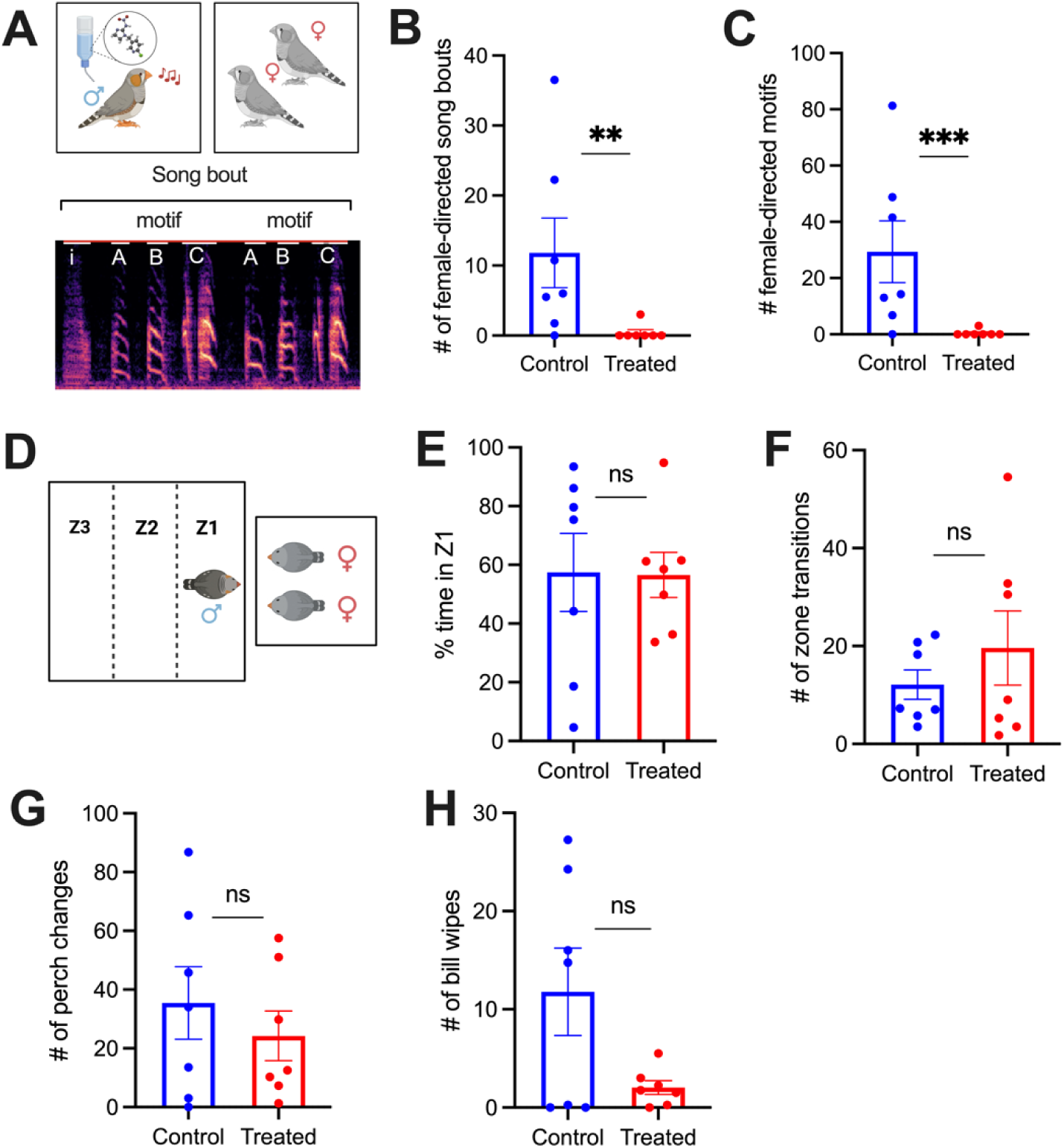
Imidacloprid exposure through drinking water suppresses courtship song without detectable changes in measured non-vocal behaviors. **(A)** Female-presentation assay and song schematic (a song bout comprising motifs). **(B**, **C)** Female-directed song bouts (**B**) and motifs (**C**) after drinking-water exposure. **(D)** Cage-zone schematic; Z1 is the zone adjacent to the female cage. **(E**–**H)** Video-scored behaviors: time in Z1 (**E**), zone transitions (**F**), perch changes (**G**) and bill wipes (**H**). Points, individual male means across four female-presentation sessions; bars, group mean ± s.e.m. Song bouts and motifs were analyzed with negative-binomial generalized linear mixed models (song bouts: χ²₁ = 10.14, *P* = 0.0014; motifs: χ²₁ = 10.99, *P* = 0.00091); non-vocal behaviors with beta (time in Z1) or negative-binomial (count) models (all Benjamini–Hochberg-adjusted *P* = 0.96). Control males sang in 21 of 28 presentations and treated males in 3 of 28 (all from one male). Control, blue; imidacloprid-treated, red. *n* = 7 control and 7 imidacloprid-treated males. ns, not significant; \*\**P* < 0.01; \*\*\**P* < 0.001.

### No detectable change in measured song structure

We next asked whether imidacloprid changed song structure by analyzing undirected song motifs sampled before and after 30 days of treatment. We found no evidence for treatment-specific changes in motif duration, average entropy, 90% frequency bandwidth or center frequency (Treatment × time interactions: all *P* ≥ 0.32; **S2 Fig; S4 Table**). Likewise, the rendition-to-rendition consistency of these features within syllables, measured as their coefficient of variation, showed no detectable treatment change (all *P* ≥ 0.52; **S2 Fig, panels F–I; S4 Table**). Thus, imidacloprid exposure through drinking water reduced song quantity without detectably altering the measured temporal or spectral features of song motifs.

### Acute injection suppresses courtship song with partial non-vocal impairment

We finally tested whether acute imidacloprid suppressed female-directed song following intramuscular (pectoral) injection. In a within-subject experiment, males received vehicle or imidacloprid injections of 5% of the estimated LD₅₀ and were tested in 5-min female-presentation assays 40 min after the injection (**Fig 3A**). Acute imidacloprid injections reduced both song bouts and motifs relative to vehicle treatment (song bouts: χ²₁ = 17.31, *P* < 0.0001; motifs: χ²₁ = 20.08, *P* < 0.0001; **Fig 3B,C; S5 Table**) and increased the latency to resume undirected singing after the assay (χ²₁ = 4.30, *P* = 0.038; **Fig 3D**). Unlike drinking-water exposure, injections produced partial behavioral impairment during female-presentation assays (**Fig 3E–I; S5 Table**). After Benjamini–Hochberg adjustment across the four non-vocal outcomes, acute treatment reduced perch changes (χ²₁ = 27.15, adjusted *P* < 0.0001; **Fig 3H**), whereas time spent near females (χ²₁ = 4.06, adjusted *P* = 0.088; **Fig 3F**), zone transitions (χ²₁ = 2.43, adjusted *P* = 0.16; **Fig 3G**) and bill wipes (χ²₁ = 1.92, adjusted *P* = 0.17; **Fig 3I**) were not detectably altered. Occasional wing droop and transient loss of balance were also observed.

**Fig 3.**
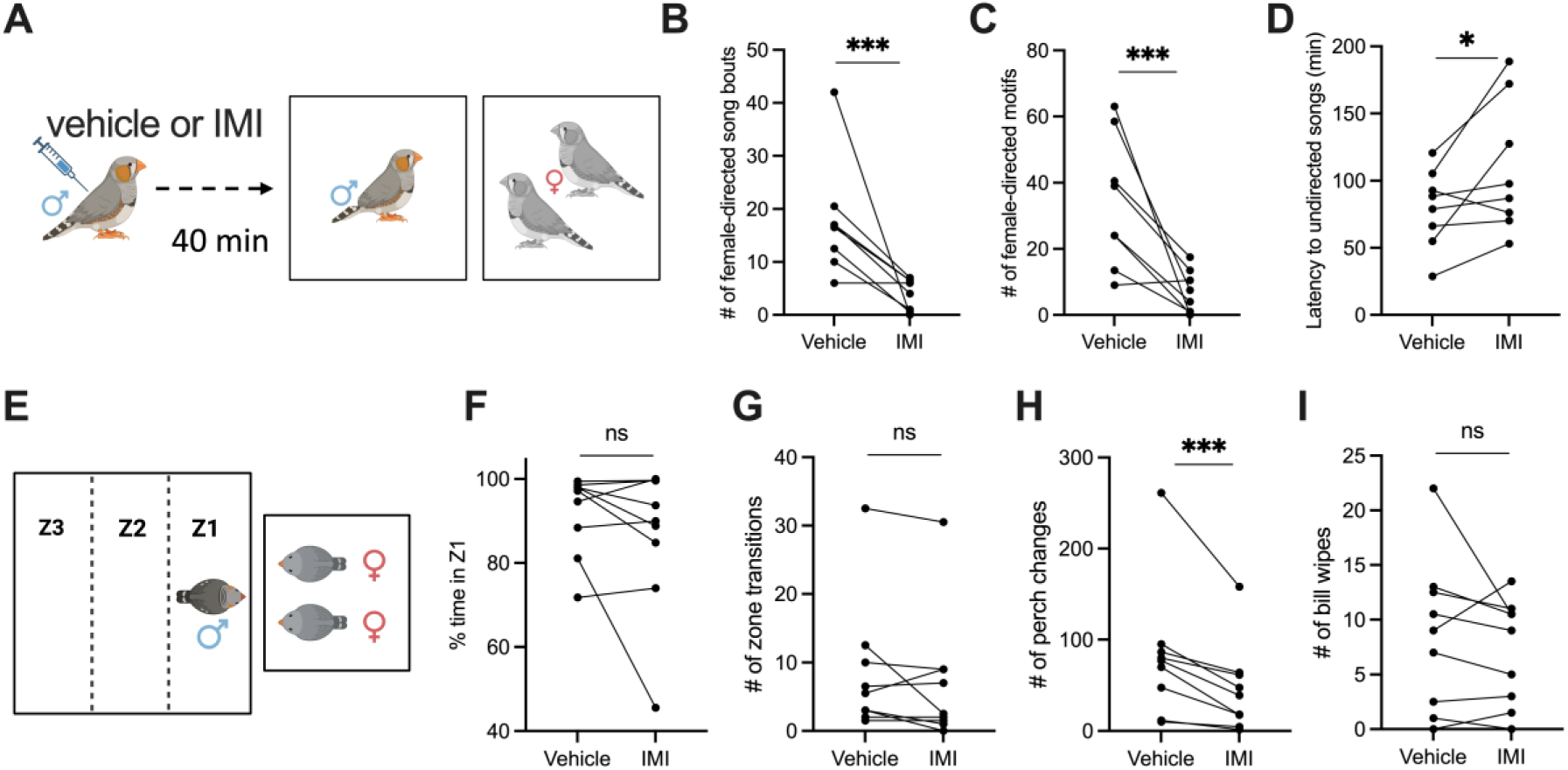
Acute imidacloprid injection suppresses courtship song and produces partial behavioral impairment during female presentation. **(A)** Acute-injection assay: vehicle or imidacloprid (5% of the estimated LD₅₀), with female presentation 40 min later. **(B**, **C)** Female-directed song bouts (**B**) and motifs (**C**). **(D)** Latency to resume undirected singing after the assay. **(E)** Cage-zone schematic; Z1 is the zone adjacent to the female cage. **(F**–**I)** Video-scored behaviors: time in Z1 (**F**), zone transitions (**G**), perch changes (**H**) and bill wipes (**I**). Points, male-level means; lines connect vehicle and imidacloprid values from the same male. Song bouts: χ²₁ = 17.31, *P* < 0.0001; motifs: χ²₁ = 20.08, *P* < 0.0001; latency: χ²₁ = 4.30, *P* = 0.038 (ratio = 1.37, 95% CI 1.03–1.83). After Benjamini–Hochberg adjustment across the four non-vocal outcomes, perch changes were reduced (χ²₁ = 27.15, adjusted *P* < 0.0001), whereas time in Z1 (χ²₁ = 4.06, adjusted *P* = 0.088), zone transitions (χ²₁ = 2.43, adjusted *P* = 0.16) and bill wipes (χ²₁ = 1.92, adjusted *P* = 0.17) were not detectably altered. Vocal analyses: *n* = 8 males, 32 sessions (one non-singing male excluded). Video analyses: *n* = 9 males, 35 sessions (one session excluded for a technical fault). ns, not significant; \**P* < 0.05; \*\*\**P* < 0.001. IMI, imidacloprid.

## Discussion

Our results show that sublethal imidacloprid exposure through drinking water and acute imidacloprid injection both suppressed song production in male zebra finches. Although previous research has established broader effects of neonicotinoids on avian physiology, reproduction, and migration, their effects on vocal communication remain largely unexplored. Drinking-water exposure reduced both undirected and female-directed singing, while acute injection also reduced female-directed song. Importantly, during drinking-water exposure, reduced song output occurred without detectable changes in body mass, female approach, measured locomotor activity, non-vocal displays, or motif structure. This dissociation suggests that song production might be disrupted without detectable changes in the broader behavioral measures examined or signs of overt toxicity. These results extend previous evidence that imidacloprid can disrupt avian reproductive physiology and behavior [11] and are consistent with an earlier descriptive report of reduced singing under pesticide exposure in red munias (*Amandava amandava*) [17].

Our findings identify singing as a potentially sensitive behavioral endpoint of neonicotinoid exposure in birds. In wild songbirds, reduced song output could compromise mate attraction, pair formation and maintenance, and territory defense, depending on the species and ecological context. Singing suppression therefore provides a plausible behavioral pathway through which sublethal pesticide exposure could reduce reproductive success. Our study did not measure reproductive outcomes or did not determine whether comparable effects occur under field exposure conditions. Nevertheless, imidacloprid exposure through drinking water suppressed a sexually and socially relevant behavior at doses that produced no detectable changes in body mass or the measured non-vocal behaviors. Neonicotinoids might therefore contribute to impaired avian reproduction not only through more direct effects on condition and survival [12–17], but also by disrupting the vocal behaviors through which reproductive opportunities are established and maintained.

Reduced singing remained detectable during the final 10 days of the 30-day washout, suggesting that the effects of imidacloprid exposure through drinking water can extend beyond the period of active exposure. If comparable persistence occurs in free-living birds, reduced singing could continue into subsequent reproductive or territorial interactions. This possibility is relevant given the detection of imidacloprid in wild birds [10] and population-level associations between neonicotinoid exposure and avian declines [6,7]. However, further studies are needed to determine whether reductions in singing occur under field exposure conditions and whether they contribute to observed avian declines.

Our finding that chronic exposure to imidacloprid suppresses singing without producing apparent effects on non-vocal behaviors, body mass, or song structure strongly suggests a potential mechanism through which low-level imidacloprid exposure selectively affects the song production system. In zebra finches, the neural mechanisms underlying song production and learning have been extensively investigated, and specialized neural circuits comprising multiple interconnected brain regions have been identified [30]. Among these regions, the sensorimotor nucleus HVC (used as a proper name) is a potential direct target of imidacloprid-induced singing suppression, as disruption of neuronal activity in HVC can lead to the cessation of singing [31]. Notably, imidacloprid is known to act on vertebrate nicotinic acetylcholine receptors (nAChRs), although its affinity for these receptors is lower than that for insect nAChRs [23, 24]. In zebra finches, nAChRs are abundantly expressed in several song-related nuclei, including HVC [28]. Thus, imidacloprid may act on HVC neurons through these receptors, disrupting neural processes involved in song initiation without affecting song structure and non-vocal behaviors. Although the present study does not identify the neural circuits through which imidacloprid reduces singing, targeted infusion of imidacloprid into HVC or other candidate nuclei involved in song production could test whether local action within these regions is sufficient to suppress song output.

More than six decades after Rachel Carson warned of a “silent spring” [1], our findings suggest that pesticide-associated silence could arise not only through the loss of birds, but also through reduced singing among exposed individuals that survive. This disruption of song production raises the possibility that widespread neonicotinoid use could contribute to population declines by compromising the ability of wild birds to attract mates, defend territories, and successfully breed. These findings underscore the importance of considering ecologically relevant behavioral changes alongside mortality and other conventional endpoints in pesticide risk assessment. By establishing the behavioral effects of imidacloprid exposure in zebra finches, our study provides a foundation for future work to identify the underlying neural and endocrine mechanisms and determine whether reduced song output affects mate attraction or reproductive success.

## Materials and methods

### Experimental animals

Twenty-three adult male and 12 adult female zebra finches (*Taeniopygia guttata*; 100–300 days post-hatch) were used in this study. Birds were either bred in our aviary or obtained from a local commercial supplier. Males were individually housed in sound-attenuating chambers under a 14:10 h light:dark cycle throughout the study. Fourteen males were randomly allocated to the control (*n* = 7) or imidacloprid-treated (*n* = 7) group for the drinking-water exposure experiment, and nine males were used in the injection experiment. Song detection was automated using the same criteria across treatments, and video recordings were scored blind to treatment. To elicit female-directed singing, eight females, presented as four pairs, were used in the drinking-water exposure experiment. Four additional females, presented as two pairs, were used in the acute-injection experiment. All birds had *ad libitum* access to water, seed and grit. Birds were acclimated to the housing conditions for at least two weeks before the study began to minimize stress-related behavioral variation. All animal care and experimental procedures were reviewed and approved by the Animal Care and Use Committee at the Korea Brain Research Institute (protocol no. IACUC-24-00040-M1).

### Imidacloprid exposure through drinking water

Imidacloprid (Sigma-Aldrich, cat. no. 37894-100MG) was used for all exposures. Because no acute oral LD₅₀ has been reported for zebra finches, LD₅₀ values from related passerine and galliform species were compiled: house sparrow (41 mg kg⁻¹), Japanese quail (31 mg kg⁻¹) and gray partridge (13.9 mg kg⁻¹) [9]. Applying the standard allometric scaling relation for pesticide toxicity, in which LD₅₀, expressed as total dose per bird, is proportional to body mass raised to a scaling factor of approximately 1.148 [32], yielded predicted zebra finch LD₅₀ values with a geometric mean of approximately 19 mg kg⁻¹. We adopted 20 mg kg⁻¹ as a rounded estimate to define sublethal exposure levels.

A fresh imidacloprid stock solution was prepared weekly (0.40 mg ml⁻¹) by dissolving 8 mg imidacloprid in 200 µl DMSO and bringing the final volume to 20 ml with distilled water. The stock was protected from light and stored at 4 °C for no more than 7 days to minimize degradation. Treatment solutions were prepared by diluting the stock in drinking water. Daily water intake was assumed to average 4 ml per bird based on reported freshwater intake in zebra finches maintained under laboratory conditions [33]. Birds had continuous *ad libitum* access to their assigned drinking water. Imidacloprid concentrations were adjusted individually according to each bird’s body mass and assumed daily water intake to produce nominal daily exposures corresponding to defined percentages of the estimated LD₅₀. Control birds received imidacloprid-free drinking water containing DMSO at final concentrations matching those in the corresponding treatment solutions.

The exposure timeline followed the sequence shown in **Fig 1A**. Birds were first monitored for 10 days under imidacloprid-free conditions to establish individual singing baselines. Treated birds were then exposed sequentially to nominal daily doses corresponding to 1%, 2.5% and 5% of the estimated LD₅₀, each for 10 days, whereas control birds received vehicle-matched, imidacloprid-free drinking water. These exposure levels were selected to assess the effects of repeated sublethal imidacloprid exposure through drinking water while remaining below the estimated acute toxicity threshold. Following the final exposure, plain drinking water was restored for a 30-day washout phase. Undirected singing behavior continued to be monitored during washout, and data from the final 10 days were analyzed as the late-washout phase.

### Body mass and health monitoring

Body mass was recorded on the final day of the baseline phase, on day 10 of each imidacloprid exposure phase, and on the final day of the 30-day washout phase. Throughout the drinking-water exposure experiment, birds were also monitored for overt signs of distress, including lethargy, reduced activity, ruffled plumage and altered posture. No such indicators were observed. These mass and welfare assessments enabled behavioral changes to be evaluated in relation to potential signs of acute toxicity or general health decline.

### Undirected song recording

Undirected songs were recorded from birds individually housed in custom sound-attenuating chambers. Each chamber contained a single recording cage measuring either 42 × 23 × 28 cm (width × depth × height; cages 1–8) or 34 × 26 × 32 cm (cages 9–14). The two cage models had similar volumes (27.0 and 28.3 l) and were distributed across groups (cages 1–8: five treated, three control; cages 9–14: two treated, four control; Fisher’s exact *P* = 0.59); including cage model as a covariate did not alter the treatment × experimental phase interaction (χ²₄ = 35.14, *P* < 0.0001).

Each recording cage was equipped with a ceiling-mounted microphone (PRO35, Audio-Technica) connected to a mixer and preamplifier (402-VLZ4, Mackie). Audio signals were digitized through an external interface (Octa-Capture UA-1010, Roland) at a sampling rate of 32 kHz with 16-bit resolution. A custom Processing-based recording program continuously monitored input from each channel and initiated recording when it detected three to four consecutive vocal elements meeting predefined amplitude, duration and inter-element-gap thresholds. Detection parameters were optimized individually to ensure reliable capture of each bird’s songs. Recordings ended automatically after a silent interval exceeding 0.5 s, producing files that typically contained a single song bout comprising one or more motif repetitions. Non-song sounds, including cage noise and isolated calls, were removed during manual curation before analysis.

Only birds with a mean baseline output of at least 300 song bouts per recorded day were included in the singing analyses. Owing to occasional technical problems with the recording computers, data were not available for every bird on every recording day; all available valid recording days were retained in the mixed-effects analyses.

### Song structure analysis

To test whether imidacloprid exposure through drinking water altered the acoustic structure of songs, we analyzed undirected song motifs from the final day of the baseline phase and the final day of exposure. We extracted acoustic features in Raven Pro 1.5 (Cornell Lab of Ornithology) [34]. For each male, we selected 20 undirected song motifs from the last baseline day and 20 motifs from the last day of exposure, yielding 560 motifs from 14 males. To standardize sampling across individuals and timepoints, motifs were selected from the first songs produced after lights-on. We used the first complete motif from different song bouts, avoiding repeated sampling from the same bout. For each motif, we extracted motif duration, average entropy, 90% frequency bandwidth and center frequency. To assess the consistency of syllable production, each motif was also segmented into its syllables, and the same four features were measured for every syllable rendition (54 male-specific syllable types across the 14 males; 2–8 per male). For each syllable type and timepoint, rendition-to-rendition variability was quantified as the coefficient of variation (CV = s.d./mean × 100) across its 20 renditions, one from each sampled motif, and CVs were averaged across each male’s syllable types.

### Female-directed song assay and behavioral scoring

After 30 days of imidacloprid exposure, each of the 14 experimental males completed four 5-min female-presentation sessions across two consecutive days, with one session in the morning and one in the afternoon. For each session, a cage (20 × 20 × 20 cm) containing a unique pair of stimulus females, one of four total pairs, was placed adjacent to the male’s recording chamber, allowing visual and acoustic contact while preventing physical interaction. The order of female pairs was counterbalanced across males and days so that every male encountered all four pairs exactly once.

From each session, we extracted all song bouts and quantified (1) the total number of female-directed song bouts and (2) the total number of motifs summed across all song bouts (**Fig 2A**). Male behavior during each session was video-recorded from above using a wide-angle camera, and videos were scored in BORIS (Behavioral Observation Research Interactive Software) [35]. For each session, we measured the proportion of time spent in three predefined cage zones relative to the female cage (Z1: adjacent; Z2: intermediate; Z3: opposite side) (**Fig 2D**), as well as the number of zone transitions. We also quantified the primary courtship-related movements reliably detectable from overhead video: perch changes and bill wipes. These metrics provided a complementary assessment of locomotor activity and courtship display behavior during female presentation.

### Acute imidacloprid injection

In an independent within-subject experiment, nine adult males each received four intramuscular injections into the pectoral muscle: two vehicle injections and two imidacloprid injections at 5% of the estimated zebra finch LD₅₀ (1.0 mg kg⁻¹). Imidacloprid was dissolved in DMSO to prepare a 10 mg ml⁻¹ stock and subsequently diluted in phosphate-buffered saline to a final concentration of 0.156 mg ml⁻¹. The vehicle contained the corresponding concentration of DMSO in phosphate-buffered saline but no imidacloprid. Injection volume was adjusted to each bird’s body mass (6.4 µl g⁻¹). Injection order was counterbalanced, and successive injections were separated by at least 72 h. Forty minutes after each injection, males were presented with a pair of adult females for 5 min. Two female pairs were used, and each male was tested with both female pairs under both treatment conditions.

Song and video recording followed the same procedures used for the female-presentation assays. We quantified female-directed song bouts and motifs during the 5-min presentation, scored non-vocal behaviors from video recordings and measured the latency to resume undirected singing after the end of the assay. One male that did not produce song in any session was excluded from song analyses because it provided no within-individual information about treatment effects. Song analyses therefore included eight males and 32 female-presentation sessions. Video analyses included all males with available behavioral recordings; one session was excluded because of a technical problem.

### Statistical analysis

Statistical analyses were performed in R 4.6.0 [36]. Count outcomes were analyzed with generalized linear models in glmmTMB, with the error distribution selected by AICc among Poisson, NB1, NB2, ZINB1 and ZINB2 candidates with identical fixed and random effects; the simpler model was retained when candidates differed by less than 2 AICc units. For undirected singing in the drinking-water experiment, models included treatment, experimental phase and their interaction as fixed effects and bird identity as a random intercept; full-day counts were analyzed with an NB1 model and first 2-h counts with a ZINB1 model. Planned baseline-referenced contrasts were calculated using emmeans with Holm-adjusted *P* values and Bonferroni-adjusted confidence intervals. To determine when treatment effects first became detectable, both models were refitted after each exposure day, comparing the baseline with all exposure days up to that day; onset was the first exposure day with a significant treatment-by-experimental phase interaction (unadjusted *P* < 0.05) in the direction of reduced singing in treated birds. Body mass and bird-level song-structure variables were analyzed using linear mixed-effects models (lme4 for body mass; glmmTMB for song structure) with the same structure (timepoint in place of experimental phase); for song structure, motif-level measurements were first averaged within bird and timepoint, and syllable CVs across syllable types within bird and timepoint; differences in baseline-to-post change and their 95% confidence intervals were estimated from bird-level change scores.

Female-directed song and video behaviors in the drinking-water experiment were analyzed using mixed models with treatment and female-pair identity (song) or trial (video) as fixed effects and bird identity as a random intercept; song bouts, motifs, zone transitions, perch changes and bill wipes were analyzed with NB1 models and time spent near the female with a beta model. *P* values for the four video behaviors in both experiments were Benjamini–Hochberg-adjusted. Proportions of time near the female were rescaled to avoid values of exactly 0 and 1 before beta regression. Acute injection data were analyzed using within-subject models with treatment and trial as fixed effects and bird identity as a fixed blocking factor: song bouts and motifs with ZINB2 models, zone transitions with an NB2 model, perch changes with an NB1 model, bill wipes with a Poisson model, time spent near the female with a beta model and latency to resume undirected singing with a log-link Gamma model. All treatment effects and interactions were tested using likelihood-ratio tests, and the adequacy of generalized linear models was evaluated using simulation-based residual diagnostics in DHARMa.

## Data and code availability

The data supporting the findings of this study and the R code used for all analyses are deposited in Zenodo and will be made publicly available upon publication.

## Author contributions

E.B.d.S., J.L., S.L. and S.K. conceived the study and reviewed and edited the manuscript. S.K., S.L. and J.L. supervised the study. S.K. and S.L. acquired funding.

E.B.d.S. collected and analyzed the data and wrote the original draft.

## Supplementary Information

**S1 Fig.**
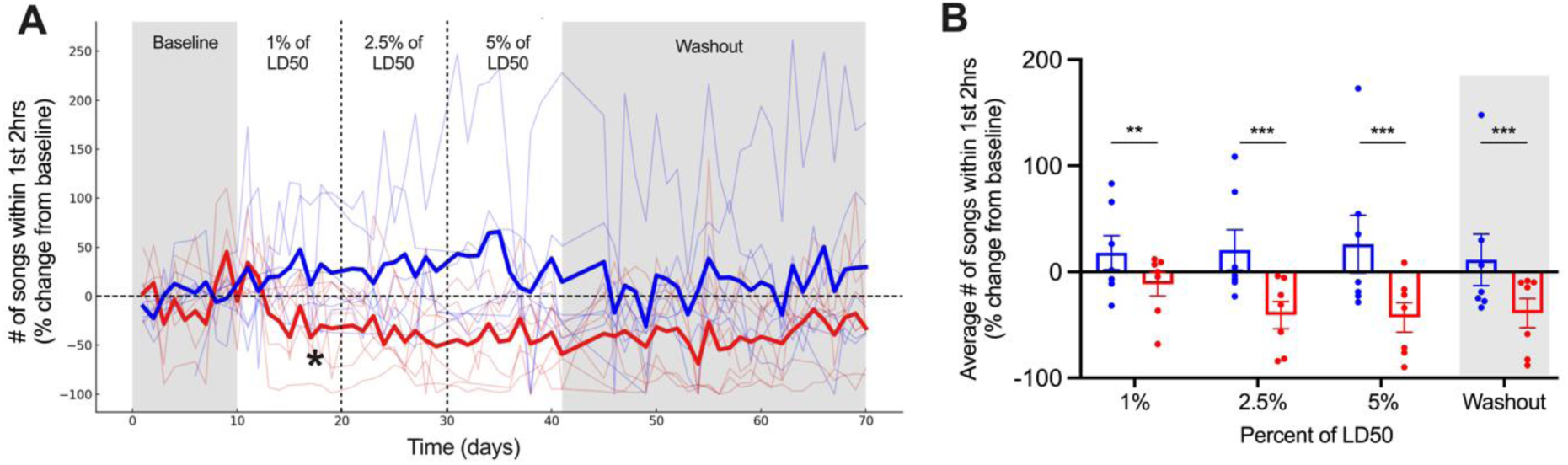
Imidacloprid exposure through drinking water reduces undirected singing during the first 2 h after lights-on. **(A)** Daily first-2-h undirected song output (% change from each male’s baseline); thin lines, individual males; thick lines, group means. The asterisk marks the first exposure day on which an exploratory cumulative analysis detected a treatment difference (seventh day of exposure; χ²₁ = 5.46, *P* = 0.019). **(B)** Mean first-2-h output during each exposure phase and late washout (% change from baseline); points, individual males; bars, group mean ± s.e.m. Analyzed with a zero-inflated negative-binomial generalized linear mixed model (treatment × experimental phase interaction: χ²₄ = 41.67, *P* < 0.0001); baseline-referenced contrasts showed reduced singing at 1%, 2.5% and 5% of the estimated LD₅₀ and during late washout. Control, blue; imidacloprid-treated, red. *n* = 7 control and 7 imidacloprid-treated males. \*\**P* < 0.01; \*\*\**P* < 0.001.

**S2 Fig.**
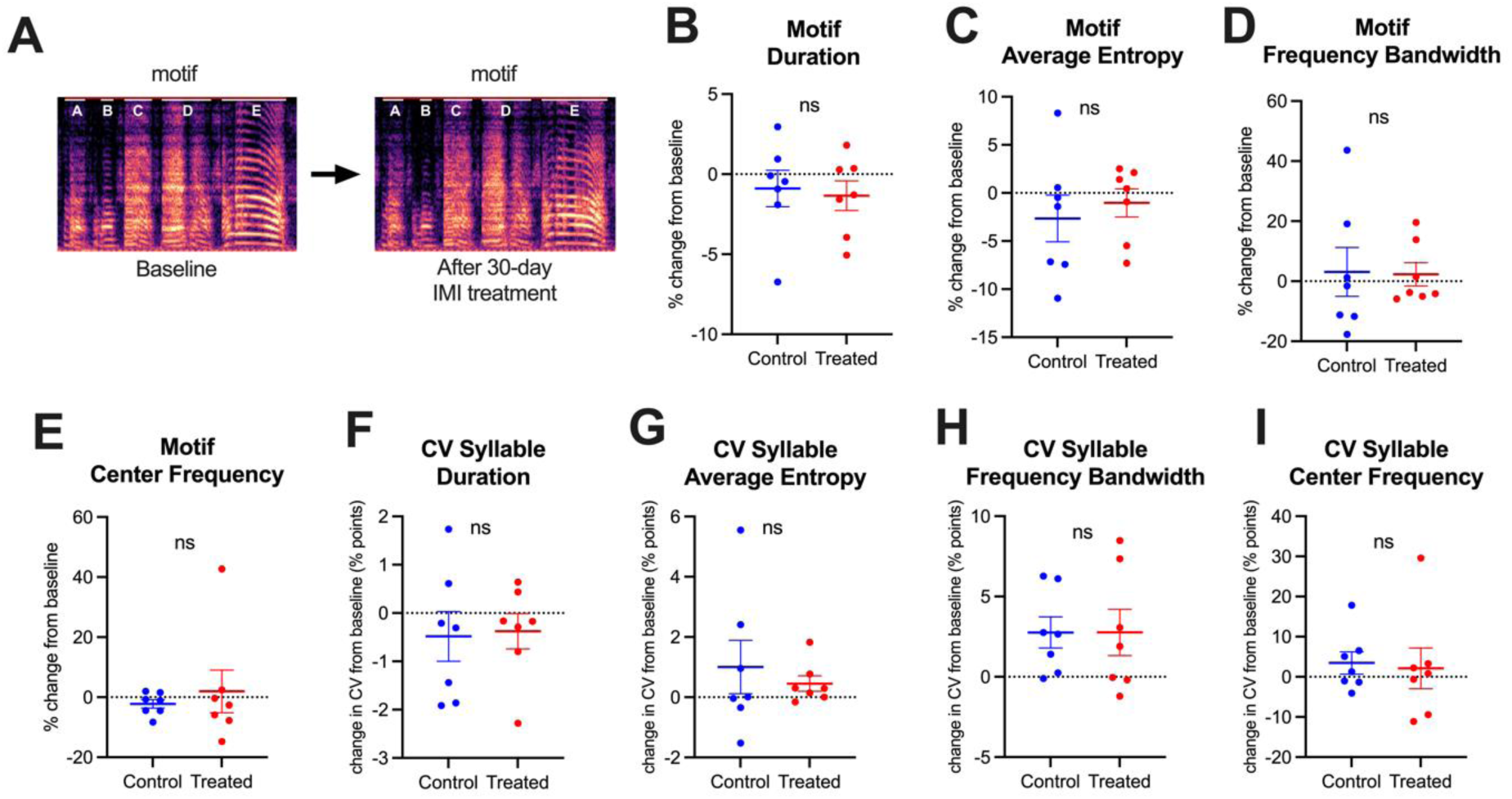
Imidacloprid exposure through drinking water does not detectably alter measured acoustic features or syllable consistency of produced songs. (A) Song-structure sampling schematic: 20 undirected motifs per male at baseline and after 30 days of exposure (560 motifs from 14 males). (B–E) Baseline-to-post change in motif duration (B), average entropy (C), 90% frequency bandwidth (D) and center frequency (E), expressed as % change from baseline. (F–I) Baseline-to-post change in the coefficient of variation (CV = s.d./mean × 100) of syllable duration (F), average entropy (G), 90% frequency bandwidth (H) and center frequency (I), expressed in percentage points; CV was calculated across the 20 renditions of each syllable type (one per sampled motif) at each timepoint and averaged across each male’s syllable types (54 syllable types; 2–8 per male). Points, bird-level changes; horizontal lines, group means ± s.e.m. Treatment × time interactions (linear mixed models) were non-significant for all features (motif duration: χ²₁ = 0.39, *P* = 0.53; average entropy: χ²₁ = 0.97, *P* = 0.32; 90% bandwidth: χ²₁ = 0.0057, *P* = 0.94; center frequency: χ²₁ = 0.89, *P* = 0.35). Treatment × time interactions for syllable CV were also non-significant (syllable duration: χ²₁ = 0.033, *P* = 0.86; average entropy: χ²₁ = 0.41, *P* = 0.52; 90% bandwidth: χ²₁ < 0.001, *P* > 0.99; center frequency: χ²₁ = 0.064, *P* = 0.80). Control, blue; imidacloprid-treated, red. *n* = 7 control and 7 imidacloprid-treated males. ns, not significant. IMI, imidacloprid.

**S1 Table.** Effects of imidacloprid exposure through drinking water on full-day and first-2-h undirected song output.

| **Outcome** | **Contrast vs baseline** | **Ratio of IRRs** | **95% CI** | ***z*** | ***P*** |
|---|---|---|---|---|---|
| Full day | 1% LD<sub>50</sub> | 1.455 | 1.128 to 1.876 | 3.685 | 0.00023 |
|  | 2.5% LD<sub>50</sub> | 1.603 | 1.247 to 2.061 | 4.693 | < 0.0001 |
|  | 5% LD<sub>50</sub> | 1.731 | 1.352 to 2.217 | 5.544 | < 0.0001 |
|  | Late washout | 1.540 | 1.200 to 1.978 | 4.315 | < 0.0001 |
| First 2 h | 1% LD<sub>50</sub> | 1.372 | 1.046 to 1.800 | 2.909 | 0.0036 |
|  | 2.5% LD<sub>50</sub> | 1.624 | 1.228 to 2.147 | 4.337 | < 0.0001 |
|  | 5% LD<sub>50</sub> | 1.982 | 1.515 to 2.594 | 6.360 | < 0.0001 |
|  | Late washout | 1.522 | 1.163 to 1.993 | 3.895 | 0.00020 |

Full-day undirected song output was analyzed with a negative-binomial (NB1) generalized linear mixed model and first-2-h output with a zero-inflated NB1 model; both included treatment, experimental phase and their interaction as fixed effects and bird identity as a random intercept (*n* = 14 males; 7 control, 7 treated). The treatment × experimental phase interaction was significant for full-day output (χ<sup>2</sup><sub>4</sub> = 35.17, *P* < 0.0001) and first-2-h output (χ<sup>2</sup><sub>4</sub> = 41.67, *P* < 0.0001). Tabulated values are ratios of incidence-rate ratios (IRRs), each comparing the control-to-treated IRR at a given phase with the corresponding ratio at baseline; values > 1 indicate that treated males sang less, relative to controls, than at baseline. Exposure phases are expressed as % LD<sub>50</sub>; late washout denotes days 61–70. *P* values are Holm-adjusted across the four planned contrasts within each outcome; 95% confidence intervals are Bonferroni-adjusted.

**S2 Table.** Body mass across imidacloprid exposure and washout.

| Timepoint | Control (g), mean ± s.d. | Treated (g), mean ± s.d. | Difference-in-differences (g) | 95% CI | *P* |
|---|---|---|---|---|---|
| Baseline | 13.47 ± 1.21 | 13.40 ± 1.29 | Reference |  |  |
| 1% LD<sub>50</sub> | 12.86 ± 0.95 | 13.17 ± 0.90 | 0.386 | −0.381 to 1.152 | 0.60 |
| 2.5% LD<sub>50</sub> | 13.04 ± 0.85 | 13.34 ± 1.01 | 0.371 | −0.395 to 1.138 | 0.60 |
| 5% LD<sub>50</sub> | 12.83 ± 0.74 | 13.07 ± 0.93 | 0.314 | −0.452 to 1.081 | 0.60 |
| Washout | 12.86 ± 0.79 | 13.27 ± 0.91 | 0.486 | −0.281 to 1.252 | 0.43 |

Values are mean ± s.d. (*n* = 7 males per treatment). Body mass was analyzed with a linear mixed-effects model with treatment, timepoint and their interaction as fixed effects and bird identity as a random intercept. The treatment × timepoint interaction was not significant (likelihood-ratio test: χ²<sub>4</sub> = 3.47, *P* = 0.48). Difference-in-differences estimates compare the treated−control difference at each timepoint with the corresponding difference at baseline; exposure phases are expressed as % LD<sub>50</sub>. 95% confidence intervals are Bonferroni-adjusted and *P* values are Holm-adjusted across the four planned contrasts.

**S3 Table.** Effects of imidacloprid exposure through drinking water on female-directed song and display behavior during female presentation.

| Outcome | Control | Treated | Effect measure | Effect estimate | 95% CI | χ² | d.f. | *P* |
|---|---|---|---|---|---|---|---|---|
| Song bouts | 11.82 ± 12.96 | 0.43 ± 1.37 | IRR (treated/control) | 0.0081 | 0.00032 to 0.205 | 10.14 | 1 | 0.0014 |
| Motifs | 29.36 ± 29.82 | 0.43 ± 1.37 | IRR (treated/control) | 0.0058 | 0.00023 to 0.146 | 10.99 | 1 | 0.00091 |
| Time in Z1 | 0.569 | 0.559 | Treated − Control (proportion) | −0.010 | −0.355 to 0.336 | 0.0030 | 1 | 0.96 |
| Zone transitions | 9.32 | 11.37 | IRR (treated/control) | 1.219 | 0.440 to 3.377 | 0.14 | 1 | 0.96 |
| Perch changes | 19.15 | 17.02 | IRR (treated/control) | 0.889 | 0.255 to 3.100 | 0.034 | 1 | 0.96 |
| Bill wipes | 3.43 | 1.23 | IRR (treated/control) | 0.359 | 0.051 to 2.505 | 0.96 | 1 | 0.96 |

Female-directed song and non-vocal display behavior were scored in the same four female-presentation sessions per male (*n* = 7 males per treatment; 28 sessions per treatment). For song bouts and motifs, Control and Treated values are session-level means ± s.d.; for the behavioral outcomes they are model-estimated means. Song-bout and motif counts were analyzed with NB1 generalized linear mixed models; zone transitions, perch changes and bill wipes with NB1 models, and time in Z1 with a beta model; all included bird identity as a random intercept. Effect estimates are treated/control incidence-rate ratios (IRRs), except time in Z1 (treated−control difference in estimated proportion). Omnibus treatment effects were tested by likelihood-ratio test (d.f. = 1); values < 1 indicate reduced output in treated birds. After Benjamini–Hochberg adjustment, all four behavioral outcomes remained non-significant (all adjusted P = 0.96). Female-pair identity did not affect song-bout (χ²<sub>3</sub> = 1.36, *P* = 0.72) or motif (χ²<sub>3</sub> = 1.37, *P* = 0.71) production.

**S4 Table.** Effects of imidacloprid exposure through drinking water on song (motif) acoustic structure and syllable consistency *Treatment × time interaction (bird-level means)*

| Outcome | Unit | Difference in baseline-to-post change | 95% CI | χ<sup>2</sup> | d.f. | *P* |
|---|---|---|---|---|---|---|
| Motif duration | s | −0.004 | −0.018 to 0.010 | 0.39 | 1 | 0.53 |
| Average entropy | bits | 0.109 | −0.147 to 0.364 | 0.97 | 1 | 0.32 |
| 90% frequency bandwidth | Hz | 24 | −712 to 759 | 0.0057 | 1 | 0.94 |
| Center frequency | Hz | 185 | −269 to 640 | 0.89 | 1 | 0.35 |
| Syllable duration CV | % points | 0.11 | −1.27 to 1.48 | 0.033 | 1 | 0.86 |
| Average entropy CV | % points | −0.55 | −2.56 to 1.46 | 0.41 | 1 | 0.52 |
| 90% frequency bandwidth CV | % points | 0.00 | −3.78 to 3.79 | < 0.001 | 1 | > 0.99 |
| Center frequency CV | % points | −1.36 | −13.92 to 11.21 | 0.064 | 1 | 0.80 |

Song-structure analysis of 560 undirected song motifs (20 per bird at baseline and 20 after 30 days of exposure). Bird-level means were the statistical unit. For each acoustic feature, a Gaussian mixed model included treatment, time and their interaction as fixed effects and bird identity as a random intercept. The upper four rows give motif acoustic features; the lower four rows give syllable consistency (coefficient of variation, CV; differences in percentage points). The difference in baseline-to-post change is the treated change score minus the control change score. There was no evidence that imidacloprid exposure through drinking water produced treatment-specific changes in the measured acoustic structure of motifs or in syllable consistency.

**S5 Table.** Acute imidacloprid injection: effects on female-directed song, post-assay latency and non-vocal behavior.

| **Outcome** | **Males** | **Sessions** | **Effect type** | **Imidacloprid / vehicle** | **95% CI (unadjusted)** | **χ²** | **d.f.** | ***P*** |
|---|---|---|---|---|---|---|---|---|
| Song bouts | 8 | 32 | IRR | 0.325 | 0.195 to 0.540 | 17.31 | 1 | < 0.0001 |
| Motifs | 8 | 32 | IRR | 0.260 | 0.155 to 0.436 | 20.08 | 1 | < 0.0001 |
| Latency to undirected singing | 8 | 32 | Latency ratio | 1.37 | 1.03 to 1.83 | 4.30 | 1 | 0.038 |
| Time in Z1 | 9 | 35 | Odds ratio | 0.583 | 0.350 to 0.971 | 4.06 | 1 | 0.088 |
| Zone transitions | 9 | 35 | IRR | 0.747 | 0.513 to 1.088 | 2.43 | 1 | 0.16 |
| Perch changes | 9 | 35 | IRR | 0.529 | 0.433 to 0.647 | 27.15 | 1 | < 0.0001 |
| Bill wipes | 9 | 35 | IRR | 0.840 | 0.656 to 1.076 | 1.92 | 1 | 0.17 |

Within-subject injection experiment (four intramuscular pectoral injections per male, two of vehicle and two of imidacloprid). Vocal outcomes were analyzed with zero-inflated negative-binomial (song, motif) and Gamma (latency) generalized linear models in the eight males that sang in at least one session (32 sessions). Video-scored non-vocal behaviors were analyzed in all nine males with usable video (35 sessions; one additional session was unscorable): time in the zone nearest the female with a beta model, zone transitions and perch changes with negative-binomial models and bill wipes with a Poisson model. All models included treatment, trial and bird identity as fixed effects, with bird identity as a blocking factor; treatment effects were tested by likelihood-ratio test (d.f. = 1). Ratios are imidacloprid-to-vehicle estimates; values < 1 indicate reduced output or odds, and values > 1 indicate longer latency. Effect estimates and 95% confidence intervals are model-based; confidence intervals are unadjusted. *P* values for the four non-vocal behaviors were Benjamini–Hochberg-adjusted; *P* values for vocal outcomes and latency were unadjusted.

